# Root phosphatase activity is coordinated with the root conservation gradient across a phosphorus gradient in a lowland tropical forest

**DOI:** 10.1101/2023.10.30.564730

**Authors:** Xavier Guilbeault-Mayers, Etienne Laliberté

**Affiliations:** Institut de recherche en biologie végétale, Département de sciences biologiques, Université de Montréal, 4101 Sherbrooke Est, Montréal, QC, Canada H1X 2B1

**Keywords:** Phosphorus availability, Organic phosphorus, Root economic space, Limiting similarity, Environmental filtering, diagnostic indicator of phosphorus availability

## Abstract

Soil phosphorus (P) is a growth-limiting nutrient in tropical ecosystems, driving diverse P-acquisition strategies among plants. Particularly, mining for inorganic P through phosphomonoesterase (PME) activity is essential, given the substantial proportion of organic P in soils. Yet the relationship between PME activity and other P-acquisition root traits remains unclear.

We measured root PME activity and commonly-measured root traits, including root diameter, specific root length (SRL), root tissue density (RTD), and nitrogen concentration ([N]) in 18 co-occurring trees across soils with varying P availability to better understand trees response to P supply.

Root [N] and RTD were inversely related, and that axis was related to soil P supply. Indeed, both traits correlated positively and negatively to PME activity, which responded strongly to P supply. Conversely, root diameter was inversely related to SRL, but this axis was not related to P supply.

Suggesting that limiting similarity influenced variation along the diameter-SRL axis, explaining high local trait diversity. Meanwhile, environmental filtering tended to impact trait values along the root [N]-RTD axis. Overall, P availability indicator traits like PME activity and root hairs only tended to be associated with these axes, highlighting limitations of these axes in describing convergent adaptations at local sites.

## Introduction

Tropical forests occur across a wide range of geological deposits (Turner & Engelbrecht, 2011). This leads to high soil nutrient heterogeneity on a large scale (Townsend et al., 2008), which might allow high tree species diversity (Wright, 2002) and high root trait variation (Ma et al., 2018). Part of this root trait variation, however, may be found within local sites since multiple trait syndromes may coexist (Chen et al., 2013; Lugli et al., 2019; Dallstream et al., 2022). High local diversity in nutrient-acquisition strategies may suggest that the root economic space (RES) dimensions defined by Bergmann et al. (2020) may be limited to describe strategies among species, rather than community-wide strategies reflecting convergent adaptation to particular environmental conditions such as soil fertility. For instance, the opposite strategies along the root collaboration gradient (i.e. high diameter and high specific root length (SRL)) (Bergmann et al., 2020) may not necessarily provide similar benefits at both ends of an environmental gradient (Laughlin et al., 2021), such as a soil fertility gradient. Instead, these opposing strategies could be functionally equivalent in terms of their consequences for plant fitness in the same local environment. Therefore, improving our knowledge on alternative phosphorus (P) acquisition strategies and those displaying functional convergence across species within communities in response to soil P availability should lead to a better understanding of tropical trees distribution across edaphic gradients.

Phosphorus availability is considered a key limiting factor for plant productivity in the majority of tropical and subtropical forests (Vitousek et al., 2010). These forests are characterized by old and strongly weathered soils that result in generally low P availability. However, tropical rainforest soils can display significant variability in P availability, depending on the nature of geological deposits (Vitousek et al., 2010; Cleveland et al., 2011; Turner et al., 2018). As a result, various morphological, chemical and physiological root adaptations to improve P acquisition are found in tropical soils (Tarafdar & Claassen, 1988; Lambers et al., 2008; Zemunik et al., 2018). For instance, the synthesis of phosphomonoesterases (PME) (that release ester-linked phosphates) is a common physiological adaptation in response to P limitation by tropical soil microbes (Turner et al., 2018) and plants (Vance et al., 2003; Marklein & Houlton, 2011; Guilbeault-Mayers et al., 2020). Only a few studies, however, have addressed how root PME activity coordinates with other morphological and chemical root adaptations across P availability gradients. Although relatively understudied, a consistent positive correlation between PME activity and SRL was observed (Ushio et al., 2015; Cabugao et al., 2021; Lugli et al., 2019, 2021; Han et al., 2022). High SRL is often associated with an ‘autonomous’ nutrient acquisition strategy, which opposes, along the collaboration gradient, a ‘collaborating’ trait syndrome involving roots of larger diameter and higher arbuscular mycorrhizal (AM) colonization (Bergmann et al., 2020). Therefore, a positive correlation between PME activity and SRL might indicate a potential trade-off between nutrient uptake beyond the root nutrient depletion zone through AM symbiosis and P mining through soil exploration involving high PME activity and high SRL. However, the response of AM colonization to P availability gradient has been proposed to be both linear (Treseder, 2004; Ma et al., 2021) and hump-shaped (Treseder & Allen, 2002), contrasting with the consistently negative linear response of PME activity to the same gradient (Allison et al., 2007; Margalef et al., 2017). This raises questions about the consistency of this trade-off along a strong soil P gradient.

Soil PME activity tends to increase with declining soil P supply (Allison et al., 2007; Margalef et al., 2017). Similarly, mycorrhizal plant species are expected to follow a general pattern whereby they rely more heavily on mycorrhizal partners for nutrient acquisition as P supply declines (Treseder, 2004; Ma et al., 2021). Phosphomonoesterase activity should, therefore, exhibit a positive correlation with root diameter rather than SRL, as both PME activity and diameter/AM colonization are expected to be high in P-depleted soil and decrease as P availability increases. Arbuscular mycorrhizal colonization response to the soil fertility gradient, however, has also been suggested to be hump-shaped (Treseder & Allen, 2002), with AM colonization being low in P-depleted soil due to nutrient limitations impacting both plant and fungal growth, as well as being low in P-rich soil owing to a decrease in plant carbon (C) allocation towards AM symbiosis. This hump-shaped pattern, when compared with the negative linear relationship between PME activity to the same gradient, suggests that the tradeoff between P uptake through AM symbiosis and P mining through roots of high PME activity might only occur within a limited range of P availability variation (i.e. from P-depleted to P-moderate soils). Furthermore, the general pattern whereby plants rely more strongly on AM colonization in P-depleted soils has been challenged by studies showing that extra-radical fungal biomass and scavenging may decline in P-depleted soils (Lambers & Teste, 2013; Teste et al., 2016). Meanwhile, given that plants could display a wide range of strategies deviating from both patterns mentioned above in P-depleted soils (Zemunik et al., 2015; Lambers et al., 2018; Wen et al., 2019; Zemunik et al., 2018), it remains unclear which functional root traits can be used predominantly as diagnostic indicators of low P availability as conclusively as PME activity (Duff et al., 1994; Margalef et al., 2017; Guilbeault-Mayers et al., 2020).

A potential explanation for the coexistence of multiple root strategies, including those related to P acquisition, within individual communities is that several different root traits or trait syndromes could be functionally equivalent in terms of nutrient acquisition and consequences on plant fitness (Marks & Lechowicz, 2006; Raven et al., 2018; Laughlin et al., 2021), at any given level of P availability. For example, in low-P soils, the greater the number of root cortical cells, the greater the probability of being colonized by AM fungi, which explains why colonization tends to be positively correlated with root diameter (Comas et al., 2014; Kong et al., 2014). On the other hand, for species with low mycorrhizal dependency, a larger root diameter could arise from the loss of metabolically active parenchyma cells in favor of the formation of metabolically inert aerenchyma and sclerenchyma cells to reduce metabolic and respiration costs and prevent roots from collapsing, respectively (Fan et al., 2003; Ryser, 2006). Meanwhile, P released from parenchyma cells during aerenchyma formation could facilitate meeting the P requirements of newly produced fine roots, reducing nutrient cost associated to P nutrition (Fan et al., 2003). Alternatively, roots with a smaller diameter could explore a greater soil volume by reducing biomass allocation in finer roots with high SRL values (Eissenstat, 1992), whereas coarser roots could be favored to enhance root growth pressure to compensate for the mechanical impedance of denser soils (Materechera et al., 1991). Altogether, these alternative morphological-related P uptake strategies within local sites might constitute a significant source of root trait variation. Therefore, this suggests that a portion of the root trait variation captured by the RES may be better suited to describe local assembly processes rather than specific patterns of trait convergence across fertility gradients.

Inconsistent root diameter responses to nutrient gradients in tropical ecosystems support that variations in morphological P-foraging traits might be more related to within-community than across community variation. For example, Yavitt et al. (2011) reported a negative correlation between fine root diameter and soil fertility, whereas others found the opposite pattern (Zangaro et al., 2008; Ushio et al., 2015; Lugli et al., 2021). Meanwhile, Wurzburger and Wright (2015) found no significant variation in root diameter in a P and nitrogen (N) fertilization study. These contradictory results may suggest that convergent adaptations in trait values along the root collaboration gradient (i.e. diameter - SRL axis) are unlikely at local sites along a P availability gradient. Rather, it is possible that variation along the diameter - SRL axis may be unrelated to the major changes in soil fertility, including P availability, if alternative P-acquisition root strategies are functionally equivalent in terms of P acquisition. If so, PME activity may be more consistently linked to traits that enhance the stabilization of organic matter in the rhizosphere (Hallett et al., 2022), extend the size of the root’s influence zone, and increase surface area (Lynch & Ho, 2005), such as root hairs, in order to improve P mining efficiency. Meanwhile, both PME activity and root hairs may correlate positively and more strongly with traits favoring rapid metabolic activity along the other dimension of the RES, known as the conservation gradient (i.e. root N concentration ([N]) – root tissue density (RTD) axis) (Bergmann et al., 2020), such as high root [N], to ensure fast P acquisition (Freschet et al., 2021). Overall, the RES dimensions (i.e. conservation gradient and collaboration gradient) were identified using a global database (Bergmann et al., 2020), but which does not consider the influence of environmental factors on trait distribution and the relative impact of a given root strategy on plant fitness under specific environmental conditions, such as soil P supply levels. Clarifying which dimension is more closely related to soil P variation would, therefore, require studies conducted on the most abundant species along a strong P availability gradient.

To evaluate how inorganic and P-mining acquisition traits coordinate and their relation to soil P availability, our study adds key root traits of the RES and less commonly measured traits such as root branching intensity, root hair length and density to previously measured PME activity on the most abundant co-occurring trees across sites in central Panama differing strongly in total and exchangeable P (Guilbeault-Mayers et al., 2020). Our objectives were to determine (i) how soil P availability gradient is related to the root collaboration and conservation gradient (Bergmann et al., 2020) and (ii) how PME activity is related to the key fine root traits defining the RES and (iii) which root traits represent key diagnostic indicators of P availability, as opposed to root traits that vary strongly within local communities. We predicted that AM colonization would display no difference between site of contrasting P availability, since AM symbiosis benefits have been suggested to be low in both P-depleted and P-rich soils (Treseder & Allen, 2002; Lambers & Teste, 2013). Instead, we hypothesized that variation in P availability should impact root trait distribution, with low-P soil being associated with roots having high [N], thereby reflecting higher PME activity and/or rapid P acquisition as a strategy to compete microorganisms (Liu et al., 2010; Freschet et al., 2017). Given that root [N] is negatively correlated with RTD along the conservation gradient (Bergmann et al., 2020), following this hypothesis, we hypothesized that low-P soils should be associated with roots of low RTD, since low-P supply should not favor long-lived roots with high maintenance costs (Laliberté et al., 2015). We further predicted that given the high ratio between P acquisition capacity of root hairs and C cost related to their production and maintenance (Bates & Lynch, 2000; Jungk, 2001), root hair length and density should increase significantly at low P supply. Overall, our study aimed to improve our understanding of P-acquisition strategies deployed by co-occurring tropical tree species, focusing on P acquisition traits that may represent diagnostic indicators of fertility, in order to better explain tropical tree species distributions across edaphic gradients.

## Methods

### Study site

We sampled fine roots from October to December 2017 on selected tree species within six 1-ha plots that are part of a tropical rainforest plot network in central Panama (Table S1; Condit et al., 2013; Guilbeault-Mayers et al., 2020). Fine roots of the 18 most abundant tree species, determined by the highest cumulative stem basal area, were sampled in three plots at each site, resulting in a total of 45 aggregate samples at the individual scale (Table S1). If individuals of one of the most abundant species were very difficult to sample or not accessible (e.g. root systems cluttered with dead trees), the next most abundant species was sampled. Sampling effort was similar between individuals of different species, however, sampling fine roots on large individuals was more time consuming, resulting in an unequal number of individuals per species (Table S1). Plots were selected to cover the widest possible range of soil P availability across the plot network in central Panama (Condit et al., 2013; Turner et al., 2018). Three plots were classified as ‘low P’, with an exchangeable [P] (resin P) below 2 mg P kg^-1^ ranging from 0.77 to 1.39 mg P kg^-1^, while the other three plots were classified as ‘high P’, with an exchangeable [P] well above 2 mg P kg^-1^ (Table S1). The 2 mg P kg^-1^ threshold used to distinguish between low- and high-P sites is ecologically relevant as it has been shown to be able to separate plant communities with low and high affinity to soil [P] (Condit et al., 2013; Sheldrake et al., 2017; Turner et al., 2018). Elevation varied from 30 to 180 m above sea level, while annual rainfall and soil [N] (both total and inorganic) did not differ significantly between low-P and high-P sites (Table S1).

### Root PME activity assay

Phosphomonoesterase activity was measured on four root tips individually (mean length = 3.94 mm) for each individual tree in each plot following two rinses with deionized water. Root PME activity was determined using 4-methylumbelliferyl phosphate as a phosphomonoester analog. Root tips were incubated in 100 μl of 50 mM acetate buffer solution adjusted to pH 5 and 100 μl of 100 μM substrate for 30 minutes at 30 °C. The reaction was terminated by the addition of 50 μl of 0.1 M modified universal buffer (Tabatabai, 1994) at pH 12. Fluorescence of 4-methylumbelliferone was determined on a Fluostar Optima (BMG Labtech, Ortenberg, Germany) with excitation at 360 nm and emission at 460 nm. Individual roots tips were weighed after drying at 60 °C for seven days. Following this measurement, root tips were hydrated in deionized water for 30 minutes and then digitized to obtain root length using WinRhizo Pro software (Regent Instruments Inc, Québec, Canada). Further details of root PME assay methods can be found in our previous study at the same sites (Guilbeault-Mayers et al., 2020).

### Root functional traits

Root functional traits were measured on fine roots which were distinguished using a root order-based threshold (McCormack et al., 2015). For each species, the threshold was determined using morphological characteristics (i.e. variation in color, texture, diameter and rigidity). This typically resulted in the sampling of the first 2-3 root orders. All root traits were measured on these root orders, with the exception of PME activity in root tips, which was previously measured on the first root order (Guilbeault-Mayers et al., 2020). Specific root length, root diameter, root branching intensity and RTD were obtained using WinRhizo Pro software (Régent Instruments Inc, Quebec, Canada). Dry mass measurement was obtained by drying fine roots at 60 °C for 72 h to complete the measurement of SRL and RTD. Root hair density was measured on 10 segments of approximately 1 mm across 10 randomly sampled first order roots. Root hair length was measured on one randomly sampled root hair per first order root previously sampled for root hair density, using images captured with a Zeiss Axio Imager 2 microscope (software: AxioVison, Jena, Germany). A root hair index was obtained by multiplying root hair length by root hair density, generating a trait describing root hair length per unit of first order root length, as performed by Holdaway et al. (2011). Root [N] was measured using an elemental analyzer (Vario Micro tube; Elementar, New Jersey, United States).

### Mycorrhizal colonization

Fine roots were cut into sections of approximately 1 cm long and then bleached in modified syringes (Claassen & Zasoski, 1992) using KOH (10% v/v) at 90 °C. The roots were removed from the KOH and acidified with diluted acetic acid for five minutes. After acidification, roots were stained in Sheaffer Black ink and vinegar (5% v/v acetic acid) solution for 4 minutes (Vierheilig et al., 1998). Afterwards, roots were conserved in a lactoglycerol solution for 48 hours to remove excess staining. Finally, roots were mounted on a microscope slide with glycerol.

Among all fungal structures, we specifically chose to only report arbuscules by root length (hereafter AM colonization) to avoid misdiagnosis of AM structures (Brundrett, 2009). As was reported in another study conducted in another subtropical ecosystem in Brazil (Zemunik et al., 2018), many root segments in our samples were intensively colonized by dark septate endophytes and other non-mycorrhizal hyphae with regularly septation. To obtain a quantitative estimate of AM colonization intensity, all arbuscules were counted in approximately 300 mm of root fragments per segments of roughly 0.15 mm and divided this number by total root length analyzed by individual.

### Statistical analyses

Major axes of root trait variation were assessed using principal component analysis (PCA) with the rda function from ‘vegan’ package (Oksanen et al., 2020). Pairwise Pearson correlation coefficients among root traits was determined using the rcorr function from the package ‘Hmisc’ (Harrell & Dupont, 2020). Differences in functional root traits and soil descriptors among soil P availability classes were assessed with linear models (lm), generalized linear models (glm) from the ‘stats’ package (R Core Team, 2020) and generalized least squares (gls) from the ‘nlme’ package (Pinheiro et al., 2020). Models were modified with an appropriate variance function or appropriate family distribution to minimize heteroscedasticity and maximize normality of models’ residuals and selected based on visual inspection of residual distributions and on the Akaike information criterion (AIC) (Zuur et al., 2009; Zuur et al., 2010). Post-hoc Tukey HSD tests were conducted using the ‘emmeans’ (Lenth, 2020) and ‘multcomp’ (Hothorn et al., 2008) packages. Finally, analyses were conducted directly on the raw data instead of using species trait means. Individuals of the same species often did not occur in the same plots, and when they did, their fine roots were not necessarily exposed to the same soil condition. Soil nutrient concentrations are known to vary by an order of 2 to 3 over a few centimeters (Chapin, 1980).

## Results

Principal component analysis revealed that variation in root [N] and RTD was coordinated with soil exchangeable P, such that roots of higher root [N] were associated with P-poor soils and roots of high RTD were associated with P-rich soils. Meanwhile, variation in root diameter, AM colonization and SRL was unrelated to the soil P gradient (Fig. 1). The first two principal component (PC) axes represented relatively similar amounts of root trait variation (PC1 = 39.0%, PC2 = 24.0%) (Fig. 1).

**Figure 1:**
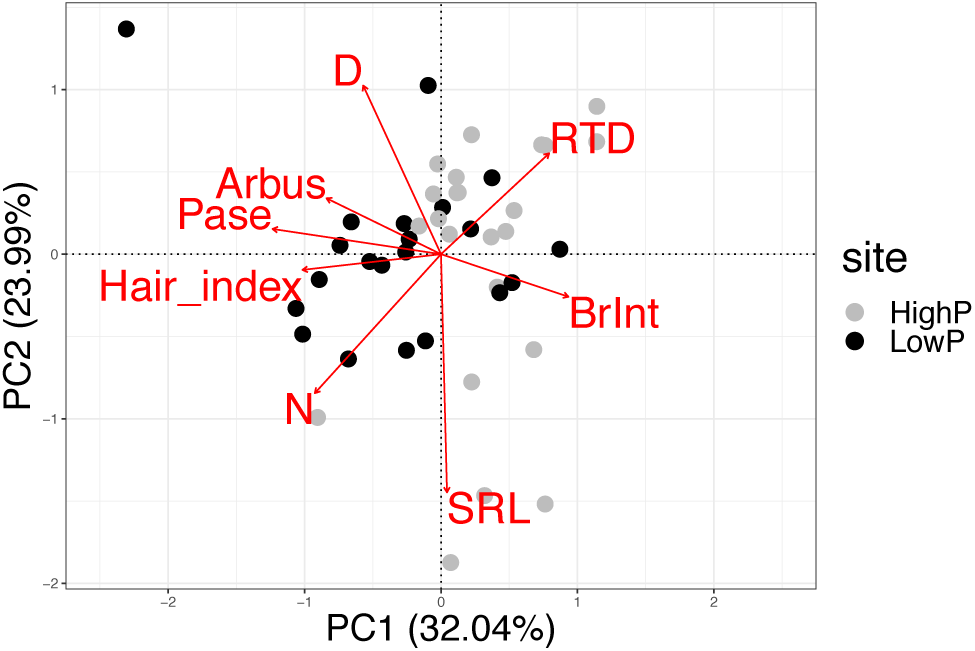
Projection of individuals in a Mahalanobis space. Abbreviations: Pase: root tips phosphomonoesterase activity, P: root phosphorus concentration, N: root nitrogen concentration, RTD: root tissue density, D: diameter, BrInt: branching intensity, Arbus: arbuscule by length and SRL: specific root length. Analysis was performed on 18 species, resulting in a total of 44 aggregate samples at the individual scale, across three plots at each contrasting exchangeable P sites.

Linear models revealed that among root traits, only three root traits differed significantly among sites. Indeed, root branching intensity was higher in P-rich sites than in P-poor sites (P = 0.015), while root tip PME activity and root hair index were greater in P-poor sites than in P-rich sites (P < 0.001 and P < 0.0001, respectively) (Fig. 2). Arbuscular mycorrhizal colonization did not differ among sites of varying soil P availability (P > 0.05) (Fig. 2) and did not differ among plots within each site (P > 0.05) (Fig. S1).

**Figure 2:**
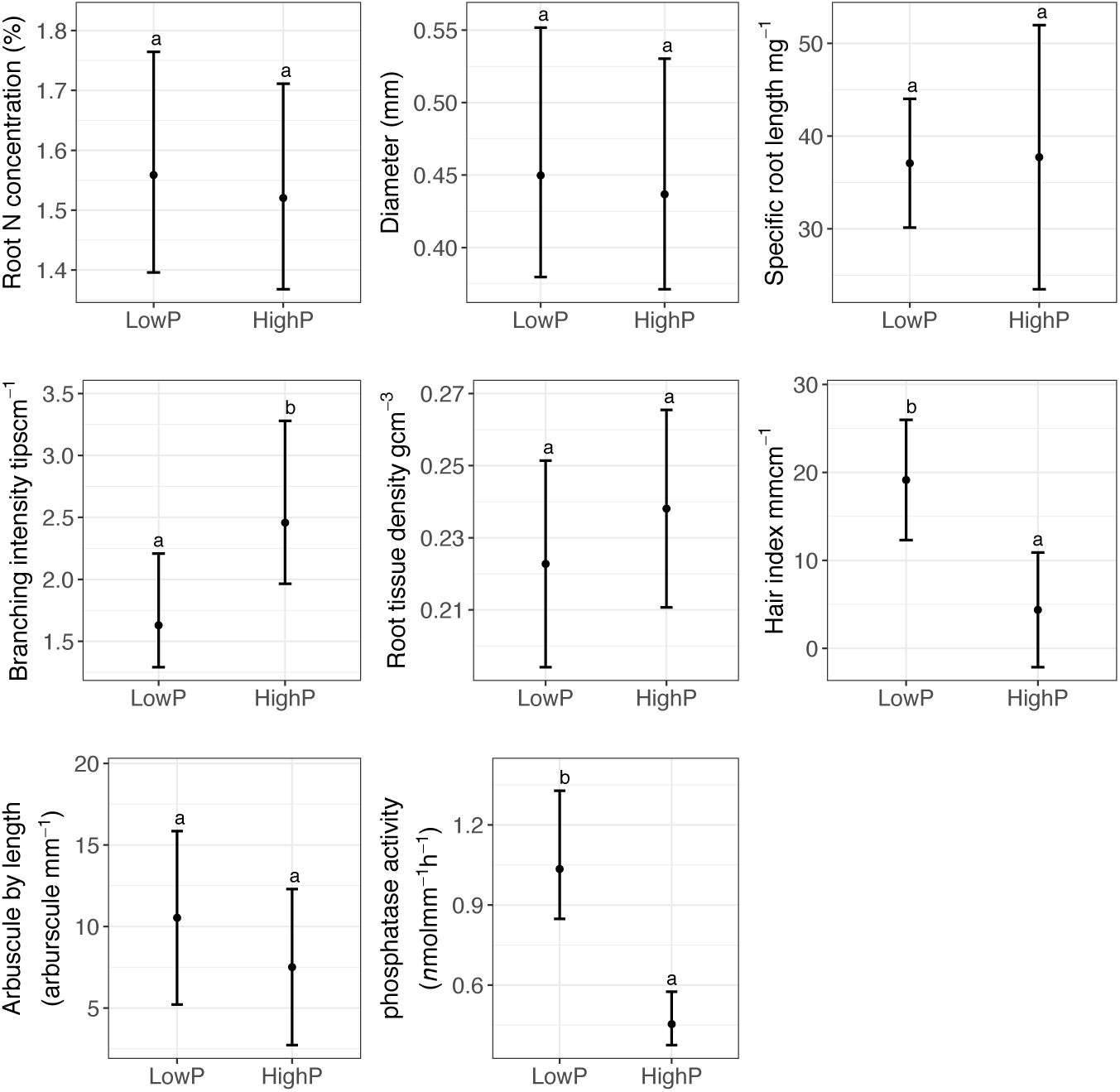
Differences in root functional traits between sites of contrasting P availability. Error bars represent the 95% confidence intervals; letters above each mean represent Tukey’s honestly significant difference (HSD) groupings (P ≤ 0.05). Mean expected values were calculated from 18 species, resulting in a total of 44 aggregate samples at the individual scale, across three plots at each contrasting exchangeable P sites.

Phosphomonoesterase activity declined with increasing RTD (*r =* −0.36, P < 0.05) and root branching intensity (*r =* −0.36, P < 0.05), while it increased with higher root [N] (*r =* 0.4, P < 0.01), root hair index (*r =* 0.45, P < 0.01), root diameter (*r =* 0.38, P < 0.05) and AM colonization (*r =* 0.32, P < 0.05). The trait distribution within the RES was confirmed, as root diameter was negatively correlated with SRL (*r =* −0.62, P < 0.00001) and RTD was negatively correlated with root [N] (*r =* −0.43, P < 0.01). However, the independence (i.e. orthogonality) of the RES dimensions was not perfect as specific root length was negatively correlated with RTD (*r* = −0.33, P < 0.05) and positively correlated with root [N] (*r* = 0.43, P < 0.01). A potential trade-off between local and long-distance soil exploration was observed since root branching intensity negatively correlated, with equal magnitude and significance, with AM colonization and root hair index (*r* = −0.31, P < 0.05). Meanwhile, AM colonization was positively correlated with the root hair index (*r* = 0.41, P < 0.01) (Fig. 3).

**Figure 3:**
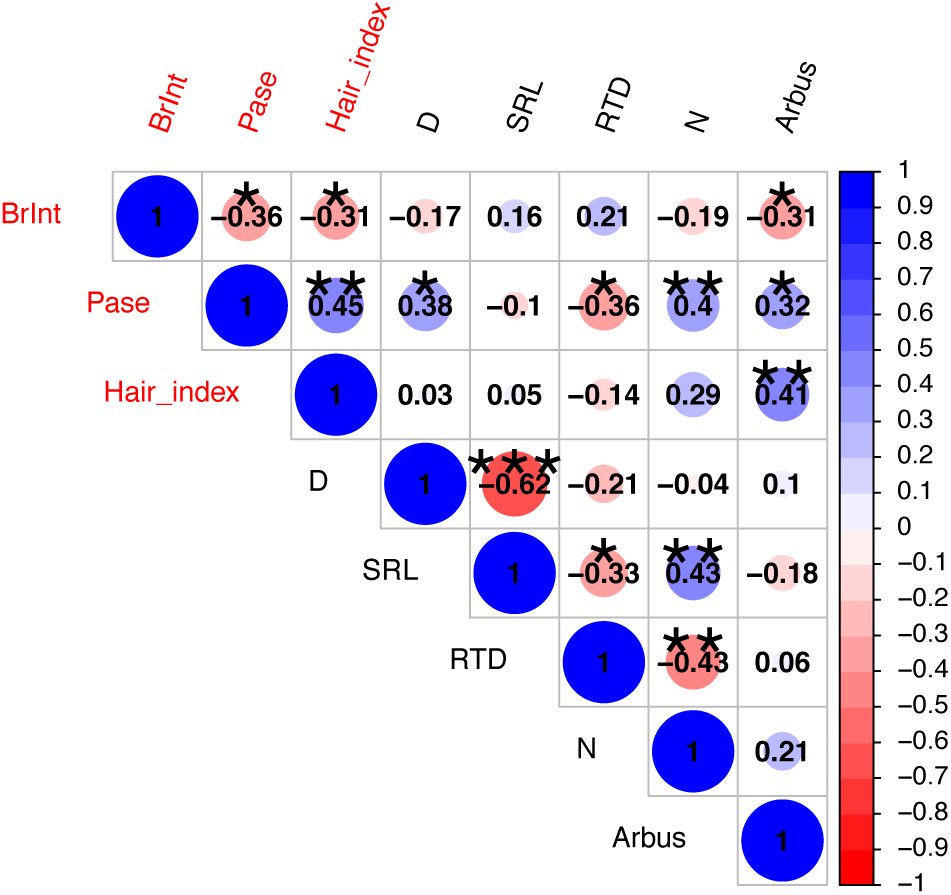
Pearson correlation among root functional traits. Stars are intended to indicate the levels of significance for four of the most commonly used levels. (*) = P < 0.05, (**) = P < 0.01, (***) = P < 0.001. Abbreviations: RTD: root tissue density, SRL: specific root length, D: diameter, Pase: root tips phosphomonoesterase activity, Hair_index: root hair length x root hair density, N: root N concentration, BrInt: branching intensity and Arbus: arbuscule by length. Root traits in red represent diagnostic indicators of P availability. Correlation coefficients were derived from 18 species, resulting in a total of 44 aggregate samples at the individual scale, across three plots at each contrasting exchangeable P sites.

## Discussion

Our results support our main hypothesis that most of the variation along the root collaboration gradient (i.e. diameter/AM - SRL axis) was observed within local sites, suggesting that different investments in AM colonization results in equivalent fitness at any given soil P level in our study. We also found evidence supporting our second hypothesis: PME activity was positively correlated with root [N] and negatively correlated with RTD, which are two traits defining the root resource conservation gradient (Bergmann et al., 2020). This provides evidence that among abundant tropical tree species, high root [N] and low RTD, two traits associated with rapid P acquisition, are likely to be favored in P-poor sites, whereas the opposite trait syndrome (i.e. low root [N] and high RTD) tends to be favored in P-rich sites. Furthermore, our third hypothesis was also supported, as PME activity positively correlated with root hair index, which promotes P acquisition through an increased surface area and the extension of the fine root influence zone. Despite this, overall, we found strong local variation in fine root traits that varied independently of soil P availability, suggesting a possible contribution of limiting similarity on plant community assembly specifically along the root diameter - SRL axis. That is, our results suggest that local competition for nutrients among co-occurring tropical tree species may lead to a higher diversity in foraging-related traits. Conversely, we also found evidence for environmental filtering for certain root traits. Specifically, root PME activity, root hair index, and branching intensity varied consistently along the soil P gradient, as well as trait values along the other dimension of the RES, the conservation gradient (i.e. root [N] - RTD axis), albeit to a lesser extent.

### Relationship between the root collaboration gradient and soil P availability

We found evidence that the traits defining the root collaboration gradient (i.e. diameter and SRL) did not vary strongly between sites of contrasted soil P availability. Most of the variation along diameter - SRL axis occurred mainly within local communities, suggesting that trait values along this axis may represent equivalently beneficial alternative P-acquisition strategies in both low and high P soils among the most abundant species. These findings align directly with prior research indicating no significant differences in root diameter (Wurzburger & Wright, 2015) and indirectly with other studies in tropical ecosystems showing either a decrease or increase in root diameter in response to P availability (Zangaro et al., 2008; Yavitt et al., 2011; Ushio et al., 2015; Lugli et al., 2021). Altogether, this points to a high local variability in root morphological adaptations associated with soil exploration (Lynch, 2019). Most contrasted strategies along the collaboration gradient (i.e. high diameter and high SRL) may have even occurred at local sites involving broader aspect of the root system architecture, such as nutrient foraging through vertical soil stratification (Lynch & Brown, 2001; Duan et al., 2020). This was particularly striking between two co-occurring species in low-P sites: *Tapirira guinanensis* developed thin and highly branched roots in the topsoil, while *Aspidosperma spruceanum* produced thick, poorly branched roots in the mineral horizon, despite displaying a similar mean AM colonization (i.e. 25.34 and 23.04 arbuscule mm^-1^, respectively) (Fig. S2).

The considerable local variability in AM fungal structures, observed in both P-poor and P-rich sites, suggests that the mean investment in the AM symbiosis might be influenced by limiting similarity, and hence, it may not be easily predicted based on soil fertility. As highlighted by our results, AM colonization did not differ among and within sites of contrasting P availability. Consequently, the mean investment in AM symbiosis did not adhere to the general pattern wherein plants typically rely more on mycorrhizal partners for nutrient acquisition in lower than higher soil P (Treseder, 2004; Ma et al., 2021), nor did it align with the hump-shaped pattern proposed by Treseder & Allen (2002). In both P-poor and P-rich sites, however, the local variability in AM symbiosis may be explained by nutrient partitioning (Turner, 2008). Under high P availability, P uptake through fine roots may entails a lower C-cost than uptake via AM hyphae (Raven et al., 2018); however, C allocation in alternative P acquisition strategies, such as fine roots alone and those associated with AM fungi, may result in similar benefits (Raven et al., 2018). This could be attributed to rhizospheric spatial and temporal soil P availability variability, such that in high-P sites, root uptake by certain species may lead to a reduction of soil P availability to a point where AM uptake may become more C-efficient (Raven et al., 2018). A similar pattern may occur in low-P sites, where a greater soil volume has to be explored to acquire exchangeable P. Arbuscular mycorrhizal hyphae incur a lower C construction cost than fine roots (Raven et al., 2018), but potentially a higher C respiration cost (Jakobsen & Rosendahl, 1990). Consequently, the varying advantages of symbiosis may have promoted soil exploration through AM hyphae for some species, enabling spatial resource partitioning among species with varying levels of dependency to mycorrhizal symbiosis (Steidinger et al., 2015). Alternatively, in low-P sites, AM colonization, a trait involved in P acquisition beyond the fine root influence zone, positively correlated with the root hair index, which facilitates P uptake through the extension of the same zone (Lynch & Ho, 2005), and with PME activity, also linked to P acquisition (Duff et al., 1994). It is conceivable, therefore, that this redundancy in P uptake-related traits may suggest that investment in AM symbiosis may have conferred additional benefits unrelated to P acquisition, such as pathogen protection (Herre et al., 2007; Wehner et al., 2010). The latter is especially relevant given our findings that low-P conditions promotes root hairs production, which may serve as entry points for soilborne pathogens (Lynch & Ho, 2005; Laliberté et al., 2015). Overall, the influence of limiting similarity on trait values along the root collaboration gradient, in our study, might provide new insights into why contrasting patterns of response of AM symbiosis to soil P availability have been proposed (e.g. Treseder & Allen, 2002; Ma et al., 2021). Regarding environmental filtering, the other pivotal process in community assembly (Götzenberger et al., 2012), it is likely to have driven values along the conservation gradient (i.e. root [N] - RTD axis), and significantly impacted root PME activity, root hair index, and branching intensity values.

### Relationship between the conservation gradient and soil P availability

We found evidence supporting that the conservation gradient (i.e. root [N] - RTD axis) was related to soil P availability, as it correlated with root PME activity which strongly responded to the soil P gradient. Root PME activity was negatively correlated with RTD, positively correlated with root [N] and not correlated to SRL. This result stands in contrast with reported positive correlations between SRL and PME activity (Lugli et al., 2019, 2021; Cabugao et al., 2021; Han et al., 2022). However, a coordinated response between SRL and PME activity (Bi et al., 2023) or general nutrient-mobilizing root exudates (i.e. total soluble organic C) is not consistent across biomes (Sun et al., 2021; Sell et al., 2022; Williams et al., 2022; Bi et al., 2023). Studies have reported that among subtropical woody species, nutrient-mobilizing root exudates (Sun et al., 2021) and PME activity (Bi et al., 2023) correlated positively and negatively with root [N] and RTD, respectively, but were independent of the diameter-SRL axis. The positive and negative correlations among PME activity and the traits defining the conservation gradient (i.e. root [N] - RTD axis) identified in these studies were substantiated by our results. However, the independence observed between PME activity and the diameter - SRL axis was not entirely upheld by our study, as PME activity positively correlated with root diameter. The trait syndrome observed in our study is similar to the positive correlation between specific root surface area and PME activity found by Ushio et al. (2015). The increase in surface area in our study was, however, mediated by a larger root diameter with lower RTD, whereas in Ushio et al. (2015), it involved an increase in root length, a decrease in root diameter, and no change in RTD. The results of Ushio et al. (2015) imply long-distance foraging for P mining, whereas our results suggest local P mining coupled with a lower RTD. The lower RTD observed in our results may, however, have allowed to explore a given soil volume with lower C investment (Ryser & Lambers, 1995), resulting in an economically advantageous trade-off for P uptake in low-P sites. Meanwhile, since higher root [N] was also part of the trait syndrome involving low RTD, greater diameter and PME activity, this may have reflected greater investment in N-rich phosphatases and maintenance of a high metabolic rate to ensure rapid acquisition of newly mineralized P (Freschet et al., 2021). However, in our study, neither of these alternative P-acquisition strategies was the result of strong trait selections, as only PME activity among the trait syndrome mentioned above may have reflected high functional convergence at local site among abundant species to which high root hair index may be included.

### Diagnostic indicators of soil P availability

Plant species can build their fine roots in different ways displaying contrasted trait values within biophysical (McCormack & Iversen, 2019) and plant-plant interaction constraints (Raven et al., 2018) to facilitate nutrient uptake. In our study, within these constraints, root PME activity, root hair index and branching intensity were the only traits that represented diagnostic indicators of P availability. Root hair length and density are recognized for expanding the size of the root’s influence zone, providing a larger exchange surface area (Lynch & Ho, 2005), and enhancing rhizospheric soil aggregation, thus stabilizing soluble organic matter (Hallett et al., 2022). In P-depleted soils, these root hair benefits could have facilitated the utilization of the organic P pool through increased PME activity, enhanced the uptake of scarce exchangeable P, and promoted the acquisition of freshly mobilized rhizospheric P. Furthermore, the association between a high root hair index and high PME activity may have enabled efficient P acquisition in species displaying lower root [N], regardless of trait value variations along the diameter - SRL axis. Enhanced soil aggregation capacity provided by root hairs may reduce nutrient leaching and, consequently, lessen the necessity for rapid uptake, which typically requires high root [N] (Freschet et al., 2021). Meanwhile, the expanded exchange surface and extended root’s influence zone rendered by root hairs may partly alleviated the need to compensate the slow P diffusion to root surfaces (Lambers et al., 2006), which generally requires soil volume exploration through morphological modifications along the diameter - SRL axis.

With regard to branching intensity, we found that it was higher in P-rich than in P-poor soils; however, its role as a diagnostic indicator of broad soil P availability remains equivocal. From Drew’s (1975) work on barley, it may be concluded that root branching intensity increases in response to high nutrient concentration, particularly in nutrient-rich patches within otherwise homogeneous, nutrient-poor soil. This trend, however, was not consistently supported by other studies involving herbaceous species (reviewed in Hodge, 2004). Branching intensity has been mostly studied in herbaceous species, as studying lateral root proliferation in mature tropical trees is challenging, limiting comparisons with our study. Nevertheless, considering that branching intensity may respond significantly to nutrient-rich patches, differences observed between soils of contrasting P-availability may result from a limited sampling effort at the individual scale that failed to capture a representative measurement of branching intensity in P-poor sites. Overall, increased branching intensity might be species-specific and more closely linked to rhizospheric nutrient concentrations than to coarse soil nutrient classifications. Nonetheless, in our study, branching intensity was negatively correlated with root hair index and AM colonization, suggesting a trade-off between local soil volume exploration in P-rich soil, and long-distance P acquisition-related traits such as an extended root’s influence zone facilitated by root hairs, or the capacity provided by AM fungi for foraging beyond this zone in P-poor soils.

## Conclusion

Our results provide new insights regarding the relationship among the RES dimensions and P availability, and into how the key processes of community assembly operate along these dimensions. These insights are in line with the combined global scale findings of Ordoñez et al. (2009), which suggest that leaf [N] is positively correlated to soil nutrient variation, and Weigelt et al. (2021), which reveal that leaf [N] is positively correlated to root [N], suggesting that environmental filtering, through varying soil nutrient supply, acts on root trait values along the conservation gradient. However, our results do not clearly indicate whether nutrient-rich or nutrient-poor soil favour high root [N], and this needs further investigation. Furthermore, since the root conservation gradient is orthogonal to the collaboration gradient (Bergmann et al., 2020), as implied by our study and Weigelt et al.’s (2021) results, the range of foraging strategies underlying the collaboration gradient might be influenced by limiting similarity and, therefore, unrelated to soil fertility variation. Overall, we found that variation along the diameter - SRL axis was greater within than between sites of contrasting soil P availability, suggesting that the range of strategies along this axis may have an equivalent impact on plant fitness, irrespective of soil P supply. Trait values along the conservation gradient were more effective in separating species occurring on soils with contrasting P availability, such that P-poor communities tended to exhibit higher root [N]. However, diagnostic indicators of P availability, such as root hairs and PME activity, were more related to P-mining strategies, showing only a tendency to be coordinated with inorganic nutrient acquisition-related trait along the RES dimensions. This suggests that variation of these trait values might be more efficient in describing local community assembly processes, rather than convergent adaptation at opposing ends of a natural P availability gradient where the relative concentration of inorganic and organic P differs greatly (Walker & Syers, 1976).

## Acknowledgments

We thank Dayana Agudo, Aleksandra Bielnicka and Julio Rodriguez from the soil laboratory of the Smithsonian Tropical Research Institute and Madeleine Trickey-Massé and Arca Arguelles-Caouette from the University of Montreal for field and laboratory assistance. Funding was provided by a Discovery Grant from Natural Sciences and Engineering Research Council of Canada (NSERC; Grant number RGPIN-2014-06106 and RGPIN-2019-04537). XGM received support from the Ernst Mayr scholarships of the Smithsonian Tropical Research Institute and from the Fonds de recherche du Québec - Nature et technologies.

## Author contributions

XGM and EL designed the project and methodology; XGM collected the data; XGM analysed the data; XGM and EL interpreted the results; XGM led the writing of the manuscript. All authors contributed to the drafts.

## Data availability

The data that support the findings of this study will be openly available upon acceptance.

