## Supporting information for "Root phosphatase activity is coordinated with the root conservation gradient across a phosphorus gradient in a lowland tropical forest"

Table S1: Number of individuals, species, mean and range soil concentrations of exchangeable phosphorus, total nitrogen, inorganic nitrogen and mean annual precipitation for each site.

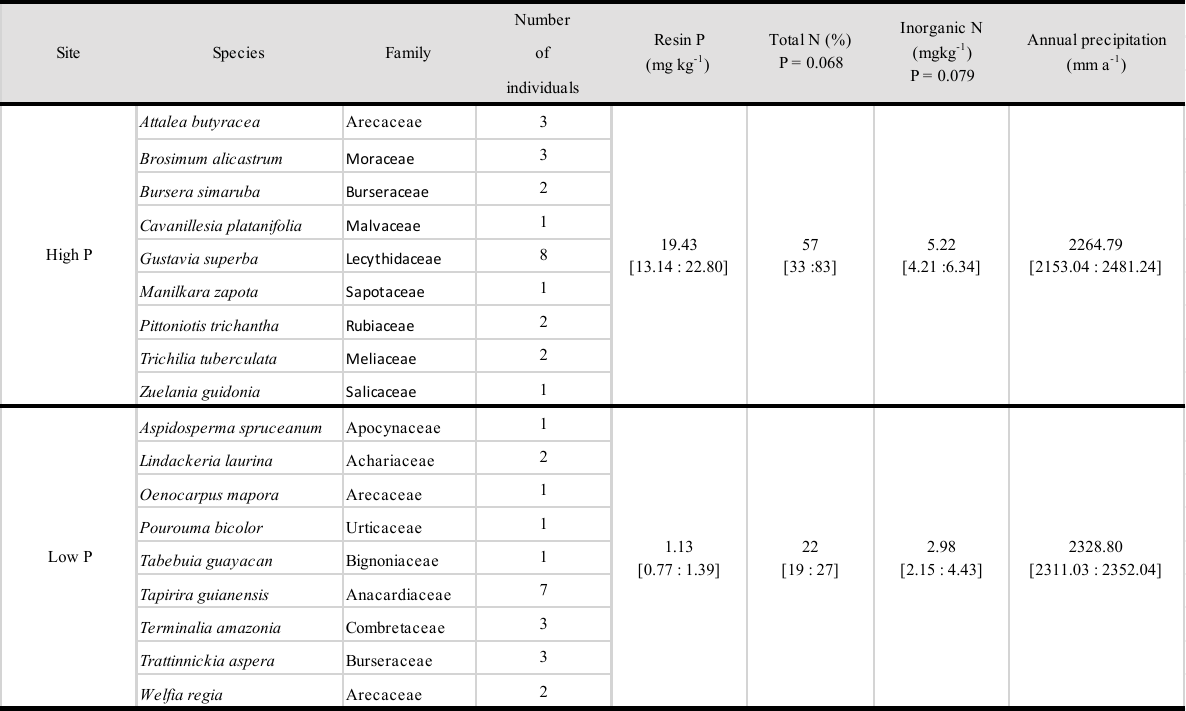

**Figure legends**

Figure S1: Differences in AM colonization among plots of contrasting P availability. Error bars represent the 95% confidence intervals; letters above each mean represent Tukey’s honestly significant difference (HSD) groupings (P ≤ 0.05). Mean expected values were calculated from 18 species, resulting in a total of 44 aggregate samples at the individual scale, across three plots at each contrasting exchangeable P sites.

Figure S2: Differences in root morphology between two co-occurring species in P-poor soil. a) *Aspidosperma spruceanum* displaying thick, poorly branched roots in the mineral horizon (between 10 and 20 cm approximatively) and b) *Tapirira guinanensis* displaying small diameter, highly branched roots in topsoil (between 1 and 5 cm approximatively). White bars represent the selection threshold for fine roots based on root orders.

Figure S1

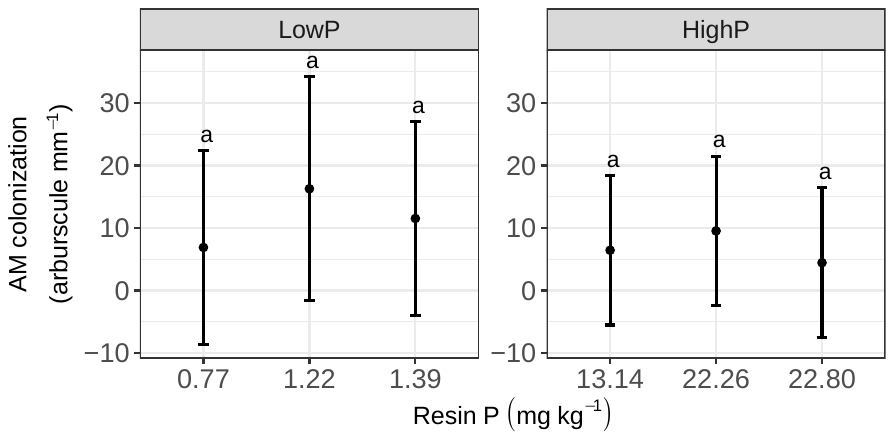

Figure S2

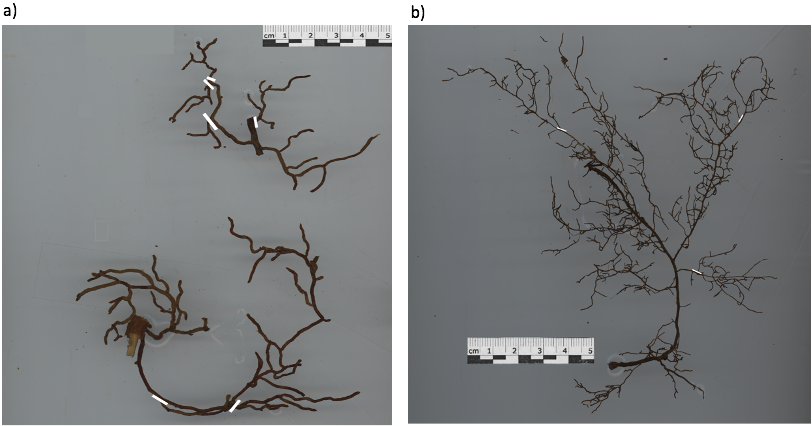
